# The yeast mitochondrial Porin represses Snf1/AMP Kinase signaling to attenuate viral replication

**DOI:** 10.64898/2026.02.25.708031

**Authors:** Sabrina Chau, Serena Marek, Aayushee Khanna, Janhavi Sathe, Sunil Laxman, Marc D. Meneghini

## Abstract

Although fungi are broadly infected with mycoviruses, the antiviral mechanisms fungal cells use to oppose viral replication are not well understood. Here we discover a new mitochondrially controlled signaling mechanism in the budding yeast *Saccharomyces cerevisiae* that limits replication of L-A, an RNA mycovirus that endemically infects this organism. We show that Por1, the mitochondrial voltage dependent anion channel, prevents hyper-replication of L-A in stationary phase cells that have exhausted media nutrients. By investigating known stationary phase regulators, we find that deletion of the AMP-activated Kinase homolog *SNF1* reverses hyper-replication of L-A observed in *por1*Δ cells. This epistatic relationship suggests that Por1 negatively regulates Snf1 in stationary phase cells and derepressed Snf1 promotes L-A hyper-replication. We confirm this model, first demonstrating that *POR1* prevents the accumulation of activated Snf1 throughout stationary phase. By investigating Snf1 signaling targets we show that this *POR1*-*SNF1* regulatory mechanism acts in stationary phase cells to limit amino acid availability that sustain L-A replication. *POR1*-*SNF1* signaling represents a novel physiological control mechanism to limit viral replication in a eukaryotic cell.

## Introduction

Recent sequencing studies have revealed a vast array of RNA viruses that infect many fungal species(*1*). Fungal mycoviruses are well known to persist as endemic infections transmitted through cell division or fusion, with no known extracellular route. Although sometimes believed to be asymptomatic, mycoviruses can profoundly affect their fungal hosts and are better understood of as spanning the symbiotic spectrum as opposed to being inconsequential travelers(*2*). For example, mycovirus infections of the Rapeseed phytopathogen *Leptosphaeria biglobosa* promote survival of the fungus through seasonal periods of high temperature that intervene crop cycles(*3*). In other cases, mycovirus infection apparently disadvantage the host as seen in phytopathogens where ‘hypoviruses’ limit the fungus’s ability to mount robust plant infections(*4*). Recent findings show that mycoviruses in human fungal pathogens elicit an opposite effect, causing hyper-virulence in cell culture and mouse infection models(*5-8*). Despite the emergence of mycoviruses as modulators of fugal pathogenesis, insights into how fungi control replication of these endosymbionts remain limited.

The L-A virus that infects *S. cerevisiae* is a comprehensively studied mycovirus, belonging to the broadly dispersed *Totiviridae* family of endogenous double stranded RNA (dsRNA) viruses. The 4.6 Kb L-A genome is encapsidated within a viral particle and extruded L-A transcripts encode for the capsid proteins (Gag) that form it. The L-A transcript also encodes a Gag-pol fusion protein that contains the canonical RNA dependent RNA polymerase (Pol) found in all RNA viruses. Each particle contains one or two Gag-pol proteins, which facilitates L-A genome replication and transcription. In many strains, L-A enables replication of satellite dsRNA segments called “M” that parasitize L-A viral particles. M satellites encode secreted toxins and their cell autonomously acting immunity, and these “Killer” toxins cause lethality in neighboring non-immune yeast(*9*).

L-A replication is maintained at a low level through the *SKI2, 3*, and *8* genes, which encode subunits of a conserved translational surveillance complex that facilitates 3’-5’ exonucleolytic RNA decay and opposes the translation of transcripts that lack polyA tails like those encoded by L-A(*10-12*). Separate pathways of L-A attenuation act through Xrn1, a 5’-3’ exoribonuclease that degrades RNAs lacking 5’ methyl caps, and Nuc1, a versatile nuclease localized to the mitochondria(*13-16*). Although L-A is asymptomatic in wild-type cells, in cells lacking *NUC1, XRN1*, and/or *SKI* genes, high levels of L-A and/or of Killer toxin cause lethality at extreme temperatures or in meiotic spore progeny(*15-18*).

Por1 is a well-known mitochondrial protein, but less well-characterized in the context of yeast antiviral responses. Earlier studies showed that L-A viral particles accumulate to high levels in *por1*Δ mutant cells that had metabolically adapted to the non-fermentable carbon glycerol for several days(*19, 20*). *POR1* encodes the yeast voltage dependent anion channel (VDAC), a highly abundant mitochondrial membrane protein. VDAC beta-barrel proteins span the outer mitochondrial membrane and control small molecule mitochondrial flux. *POR1* is essential for respiratory growth at high temperatures and plays important roles in mitochondrial osmotic stability, phospholipid metabolism, autophagy, mitochondrial protein import, and movement of phospholipids from one membrane bilayer leaflet to the other, a process known as lipid scrambling(*21-27*). In some contexts, VDACs assemble into higher order oligomers thought to facilitate movement of larger molecules, and recent findings show that Por1 lipid scramblase activity requires its oligomerized form(*26, 27*). If and how any of these functions relate to L-A repression is unknown.

Here we show that *POR1* prevents L-A from accumulating to high levels in stationary phase cells following sustained culture in standard growth media. Using genetic and biochemical experiments, we elucidate that *POR1* represses L-A by preventing hyper-activation of Snf1, the yeast homolog of AMP-activated Kinase (AMPK). By investigating Snf1 signaling, we show that this *POR1-SNF1* regulatory circuit represses L-A replication in stationary phase cells by limiting amino acids. Our findings identify a novel *POR1-SNF1* regulatory mechanism in budding yeast that may be similarly functional in other eukaryotes.

## Results

### *POR1* represses Snf1/AMPK signaling in stationary phase to prevent L-A replication

Dehanich et al found that *POR1* repressed L-A after cells had adapted to respiratory metabolism over several days(*19*). To confirm these findings in the reference S288c strain background that is naturally infected with L-A we first characterized growth and metabolism of isogenic wild-type and *por1Δ* strains through seven-day batch cultures. As expected, logarithmically growing wild-type cells vigorously fermented glucose to produce ethanol and ceased growth when glucose was exhausted (Figure 1A and 1B). While *por1Δ* displayed slower logarithmic growth and an according reduced rate of glucose consumption and ethanol production, both wild-type and *por1*Δ expended all the glucose by 20 hours of culture (Figure 1A and 1B). Over the course of the following six days both wild-type and *por1*Δ consumed all the ethanol indicating robust respiratory metabolism (Figure 1B). To investigate *POR1* repression of L-A we measured L-A Gag levels using western blots in wild type and *por1Δ* strains at intervals using these same batch culture conditions. We found a large increase of Gag levels in *por1Δ* cells by 7 days of incubation (Figure 1C). To confirm that this increase in Gag protein reflects higher viral copy number we measured L-A RNA levels using RT-qPCR, which revealed a 60-fold increase in *por1Δ* compared with wild type (Figure 1D). These findings confirm that Por1 represses L-A viral replication in respiratory cells that have exhausted nutrients. These are typically referred to as stationary phase cells, known for their metabolic quiescence and enhanced stress resistance(*28*).

**Figure 1.**
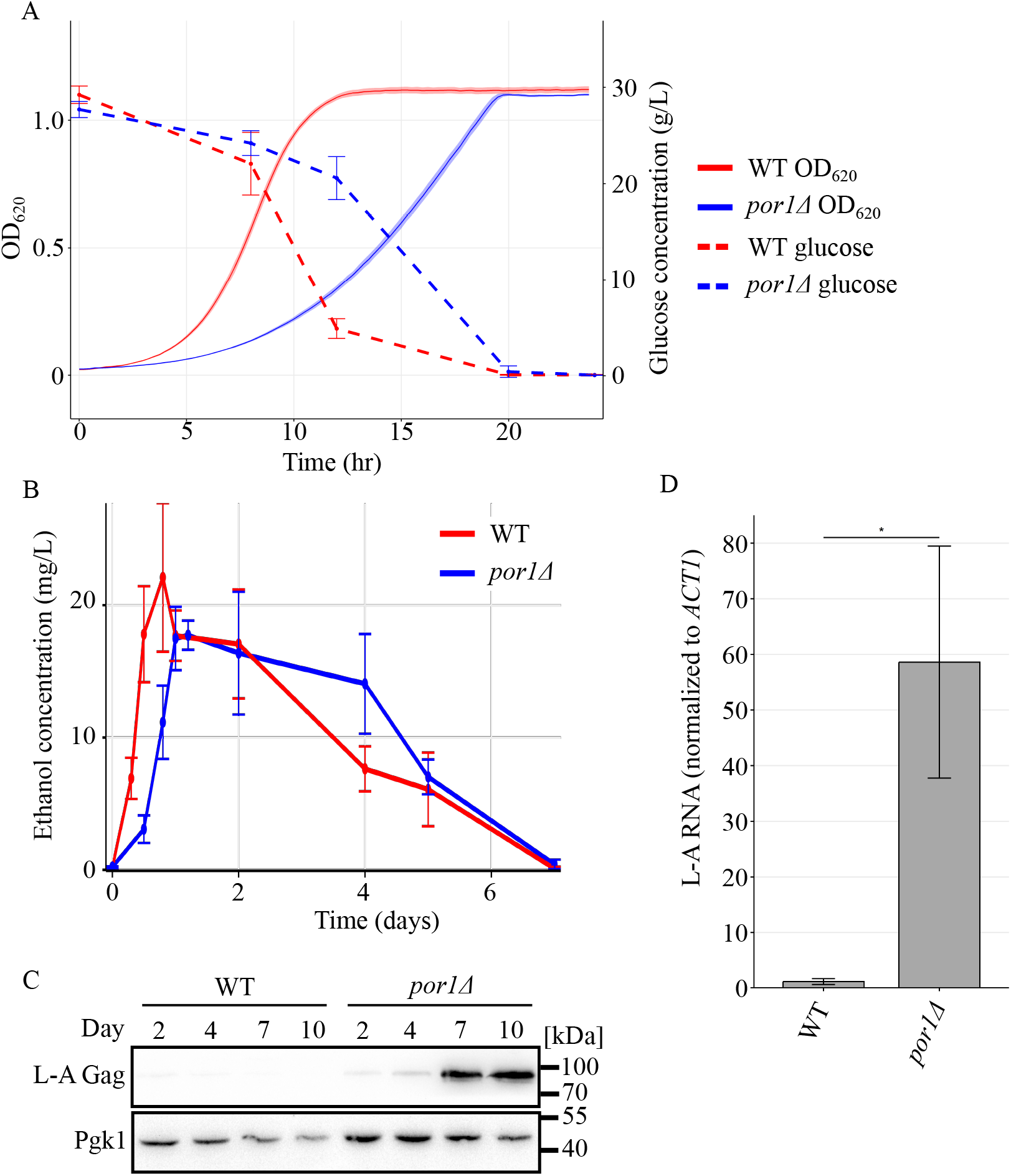
Por1 represses L-A replication after a prolonged incubation period. **(A)** Quantitative growth curve and media glucose concentration of the indicated strains. Cells were grown in YPAD at 30°C for 24 hours. Density measurements of cell cultures were taken every 15 minutes, and glucose concentration in the media was measured at indicated time points. n = 3. **(B)** Quantitative measurements of media ethanol concentration of the indicated strains. Cells were grown in YPAD at 30°C for 7 days. Ethanol concentration in the media was measured at indicated time points. n = 3. **(C)** Western blotting of L-A Gag and Pgk1 protein levels in the indicated strains. Samples were collected from cultures grown at the indicated time points in SC media supplemented with glucose. Molecular weight markers are indicated on the right. **(D)** RT-qPCR quantification of L-A RNA normalized to endogenous *ACT1* RNA. Mean RNA level and standard deviation is shown. n = 3. * *p* < 0.05. The *p* value was calculated using unpaired student’s t-test.

Progression into stationary phase involves massive gene expression reprogramming characterized by repression of nearly all transcription but activation of catabolic genes(*29*). The PAS family kinase Rim15 and the AMP-activated protein kinase Snf1 govern much of this gene expression program through phosphorylation of downstream transcriptional regulators(*30*). To investigate the roles of *SNF1* and *RIM15* for viral replication in stationary phase, we compared the levels of L-A Gag in deletion mutants of these genes by themselves or in combination with *por1*Δ. Deletion of *SNF1* completely reversed the Gag accumulation phenotype of *por1*Δ while *rim15*Δ had a comparatively minor effect (Figure 2A). This epistatic relationship suggests that Por1 represses Snf1 and that de-repressed Snf1 promotes L-A replication. To test this, we measured the accumulation of the active isoform of Snf1 phosphorylated on threonine-210 (Snf1-pT210) using western blotting experiments with an antibody that recognizes it(*31*). Supporting our model, we detected substantially higher levels of Snf1-pT210 throughout stationary phase in *por1Δ* cells compared with wild-type (Figure 2B, S1A, S1B). Moreover, Por1 levels were strongly induced in stationary phase exactly when our genetic results show that it represses Snf1-pT210 (Figure 2B, S1B). These findings demonstrate that Por1 represses Snf1 activation in post-diauxic stationary phase cells and that *SNF1* function is required for L-A hyper-activation.

**Figure 2.**
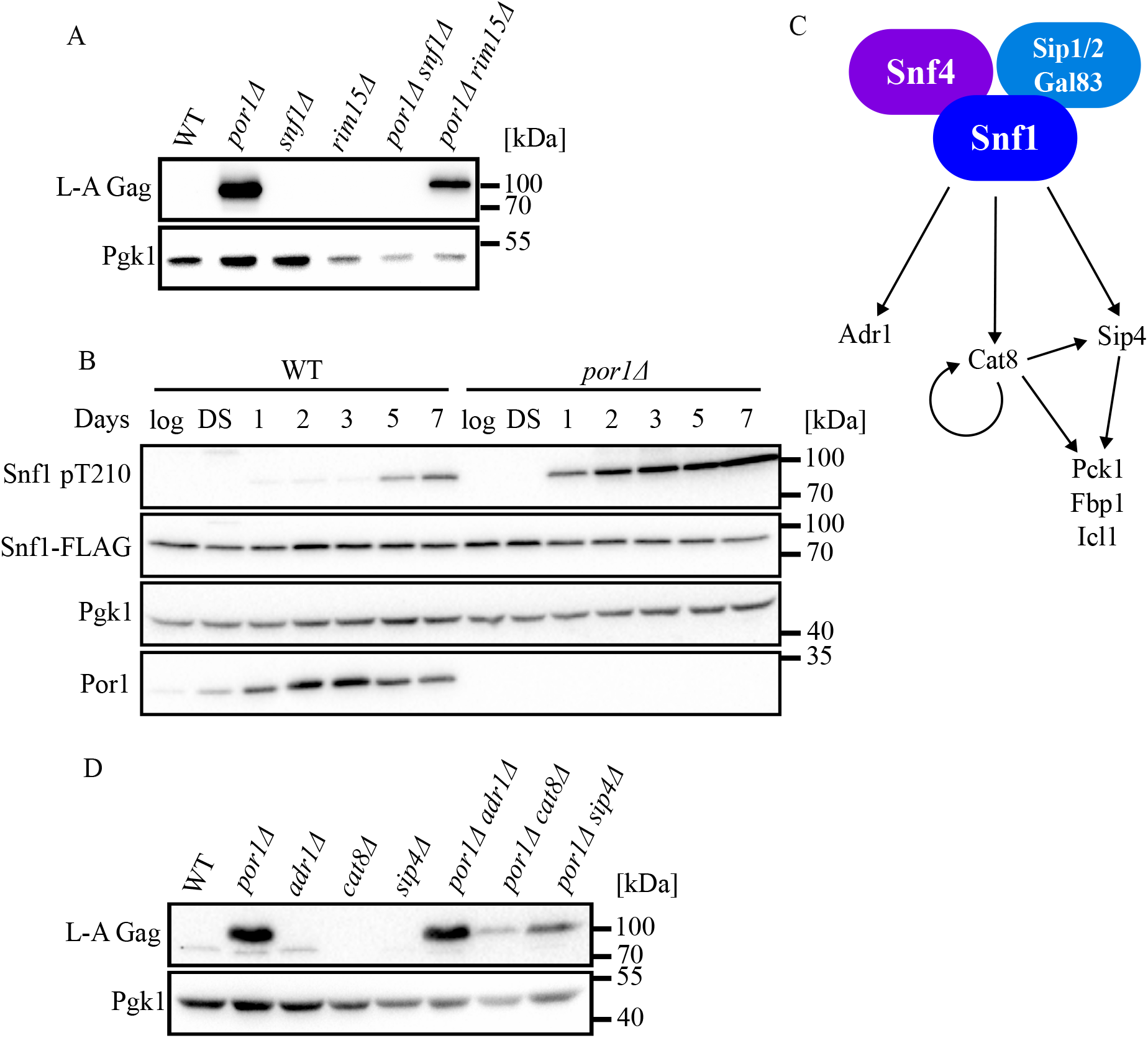
Por1 negatively regulates Snf1 activity to control L-A replication. **(A)** Western blotting of L-A Gag and Pgk1 protein levels in the indicated strains. Samples were collected from 7-day cultures grown in SC media supplemented with glucose. Molecular weight markers are indicated on the right. **(B)** Western blotting of phosphorylated Snf1, Snf1-FLAG, Pgk1, and Por1 protein levels in the indicated strains. Samples were collected from cultures grown for the indicated time points in YPAD media. Molecular weight markers are indicated on the right. **(C)** Schematic of Snf1 pathway. The Snf1 complex composes of the alpha subunit Snf1, one of the three beta subunits Sip1/2 and Gal83, and the gamma subunit Snf4. The Snf1 complex regulates expression of target genes Pck1, Fbp1, and Icl1 by phosphorylating downstream transcription factors Cat8 and Sip4. **(D)** Western blotting of L-A Gag and Pgk1 protein levels in the indicated strains. Samples were collected from 7-day cultures grown in SC media supplemented with glucose. Molecular weight markers are indicated on the right.

In addition to its activating threonine-210 phosphorylation, Snf1 is controlled through its association with co-factors in a conserved hetero-trimeric complex (Figure 2C). The gamma subunit Snf4 binds to Snf1’s autoinhibitory domain to relieve Snf1’s autoinhibition, while the beta subunits, Sip1, Sip2, and Gal83, control Snf1 activity in different subcellular localizations(*32*). We assessed the levels of L-A Gag in *por1Δ* strains lacking these proteins to discern the contribution of Snf1 subunits for viral control. Like with *snf1*Δ, *snf4*Δ completely reversed the accumulation of high Gag levels caused by *por1Δ* (Figure S1C). None of the beta subunit deletions similarly reversed the *por1*Δ phenotype, consistent with known redundancy in their control of Snf1 (Figure S1C). These findings show that Snf1 cofactors are required for its proviral function when it is hyper-activated in cells lacking *POR1*.

### Snf1 promotes L-A replication through its glyoxylate cycle targets

To elucidate how hyper-activated Snf1 promotes L-A replication we investigated its well-characterized targets, the transcription factors Adr1, Cat8, and Sip4, which drive the expression of genes involved in alternative carbon utilization in glucose starved cells (Figure 2C)(*32*). To determine if these transcription factors mediate Snf1’s proviral activity, we assessed L-A Gag levels in double mutants combining *por1*Δ with *cat8*Δ, *sip4*Δ or *adr1*Δ. While *adr1*Δ had no consequence, the high Gag levels of *por1*Δ cells were reduced when *CAT8* or *SIP4* were deleted with a marked effect caused by *cat8*Δ (Figure 2D). Cat8 and Sip4 regulate overlapping sets of genes involved in gluconeogenesis and the glyoxylate cycle (Figure 2C)(*33, 34*). To test if Cat8 and Sip4 redundantly mediate the proviral function of Snf1 we combined deletions of their shared targets *FBP1, PCK1*, and *ICL1*, with *por1*Δ. *FBP1* and *PCK1* encode gluconeogenic proteins while *ICL1* encodes a key protein of the glyoxylate cycle. We found that the high Gag levels of *por1*Δ cells were completely reverted by *icl1*Δ with only weak or no consequence of *fbp1*Δ or *pck1*Δ (Figure 3A). Collectively, these findings show that *POR1* prevents hyperactivation of Snf1 kinase in stationary phase cells, and that activated Snf1 promotes L-A replication in a manner dependent on its downstream target gene *ICL1*.

**Figure 3.**
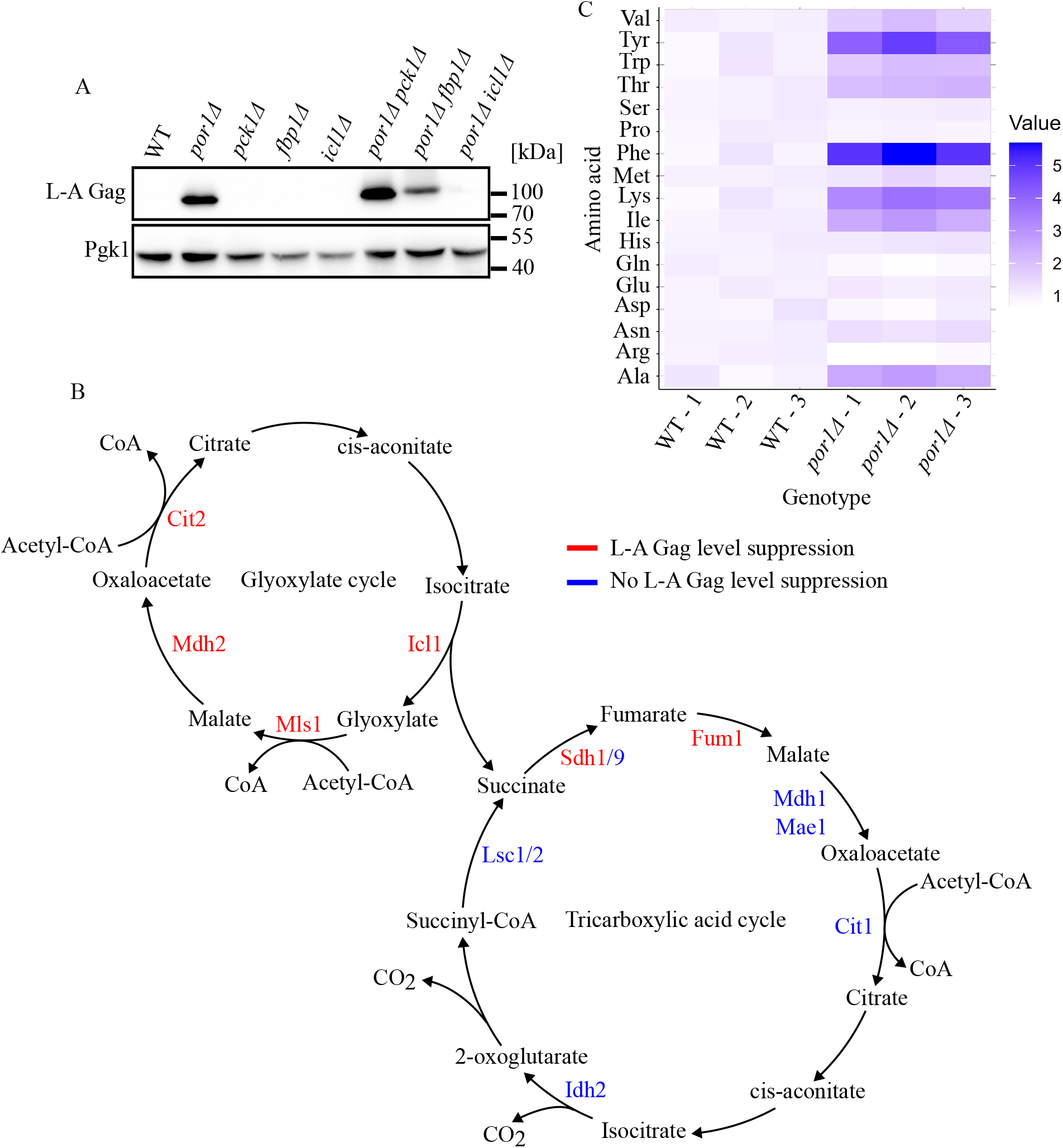
L-A replication in *por1Δ* stationary phase requires the anaplerotic glyoxylate cycle. **(A)** Western blotting of L-A Gag and Pgk1 protein levels in the indicated strains. Samples were collected from 7-day cultures grown in SC media supplemented with glucose. Molecular weight markers are indicated on the right. **(B)** Schematic of the glyoxylate and TCA cycles. The intermediates and flow of carbon through both cycles are indicated. Enzymes tested to be required for viral replication are shown in red, while enzymes dispensable for viral replication are shown in blue. **(C)** Heat map of normalized fold change amino acid levels. Prototrophic strains of the indicated genotype were used for the metabolic studies. Samples were collected from 24-hour cultures grown in SC media supplemented with glucose. Amino acid levels were measured with LC-MS/MS and were normalized to wild type. n = 3.

Icl1 catalyzes the conversion of isocitrate to succinate and glyoxylate, a key intermediate of a TCA-cycle shunt known as the glyoxylate cycle found in bacteria, fungi, plants, and some invertebrates (Figure 3B)(*35, 36*). The requirement of *ICL1* for L-A hyper-replication thus implicates the glyoxylate cycle in this process. We tested this through double mutant analysis combining *por1*Δ with other glyoxylate cycle gene deletions. As with *icl1*Δ, we found that *mls1*Δ, *mdh2*Δ, or *cit2*Δ reversed the *por1*Δ hyper-L-A phenotype (Figure 3B; Figure S2A; Table S1). While Icl1, Mls1, Mdh2, and Cit2 all act in the cytosol, glyoxylate metabolism also occurs within peroxisomes through the Mls1 and Mdh2 homologs, Dal7 and Mdh3(*37*). Deletion of *DAL7* or *MDH3* did not reverse L-A hyper-replication caused by *por1*Δ. *MDH2* and *MDH3* encode malate dehydrogenases, and a third, Mdh1, localizes to the mitochondria where it functions in the TCA cycle. Deletion of *mdh1*Δ similarly failed to revert the *por1*Δ phenotype (Figure 3B; Figure S2B; Table S1). These findings show that genes encoding cytosolic glyoxylate cycle proteins are crucial for L-A hyper-replication in *por1*Δ cells.

Succinate produced by the glyoxylate cycle can be funneled to mitochondria for TCA metabolism (Figure 3B). As the glyoxylate and TCA cycles are thus intertwined, we further tested the role of TCA cycle genes for L-A Gag accumulation caused by *por1*Δ. Like with *mdh1*Δ, deletion of most TCA cycle genes failed to revert the *por1*Δ phenotype (Figure 3B, S2B, S2C; Table S1). However, the TCA cycle mutants *fum1Δ* and *sdh1Δ* did revert *por1Δ* (Figure 3B, S2B, S2C; Table S1). As Fum1 and Sdh1 act early in the TCA cycle to metabolize succinate, a plausible explanation for these findings may be through succinate buildup causing inhibition of Icl1 and/or other glyoxylate cycle enzymes(*38*). These findings refine a model in which Por1 repression of Snf1 prevents glyoxylate cycle activation that promotes L-A replication.

### The *POR1*-*SNF1* regulatory system prevents L-A hyper-replication by limiting amino acid availability

How might the glyoxylate cycle promote L-A replication in *por1*Δ cells? The glyoxylate cycle is a versatile pathway that allows effective utilization of two-carbon units in the absence of glucose. Through this, cells obtain metabolic intermediates that support gluconeogenesis and can also be used to synthesize amino acids. An increased glyoxylate cycle will lead to more oxaloacetate, which can directly be converted to aspartic acid (which in turn supports the biosynthesis of multiple amino acids, as well as sustaining alpha-ketoglutarate production leading to glutamate/glutamine synthesis). We hypothesized therefore that enhanced amino acid availability caused by increased glyoxylate cycle flux may fuel L-A replication in *por1*Δ stationary phase cells. To test this, we first used LC-MS/MS to compare the levels of amino acids in post-diauxic wild type and *por1Δ* strains 24 hours post-inoculation. Consistent with our hypothesis, we observed substantially increased steady-state pools of multiple amino acids in *por1*Δ (Figure 3C).

We further investigated amino acid synthesis enzymes that utilize glyoxylate cycle produced precursors. Glyoxylate can be converted to glycine by the alanine-glyoxylate aminotransferase Agx1 while oxaloacetate is converted to aspartate through the mitochondrial and cytosolic localized aspartate aminotransferases Aat1 and Aat2, respectively (Figure 4A)(*39-41*). In contrast to glycine, aspartate is a versatile amino acid that can be used to produce numerous additional amino acids (Figure 4A). To test if glyoxylate cycle intermediates lead to amino acid synthesis that enhance L-A replication, we combined *por1*Δ with deletions of *AAT1, AAT2*, and *AGX1* to test for reversion of the *por1*Δ phenotype. While deletion of *AAT1* or *AGX1* had little effect on the levels of L-A Gag, *aat2*Δ reverted the accumulation of Gag in *por1Δ* (Figure 4A, B). Like the required glyoxylate cycle proteins, Aat2 is cytosolic, suggesting that L-A replication in stationary phase requires the synthesis of aspartate from oxaloacetate in the cytosol.

**Figure 4.**
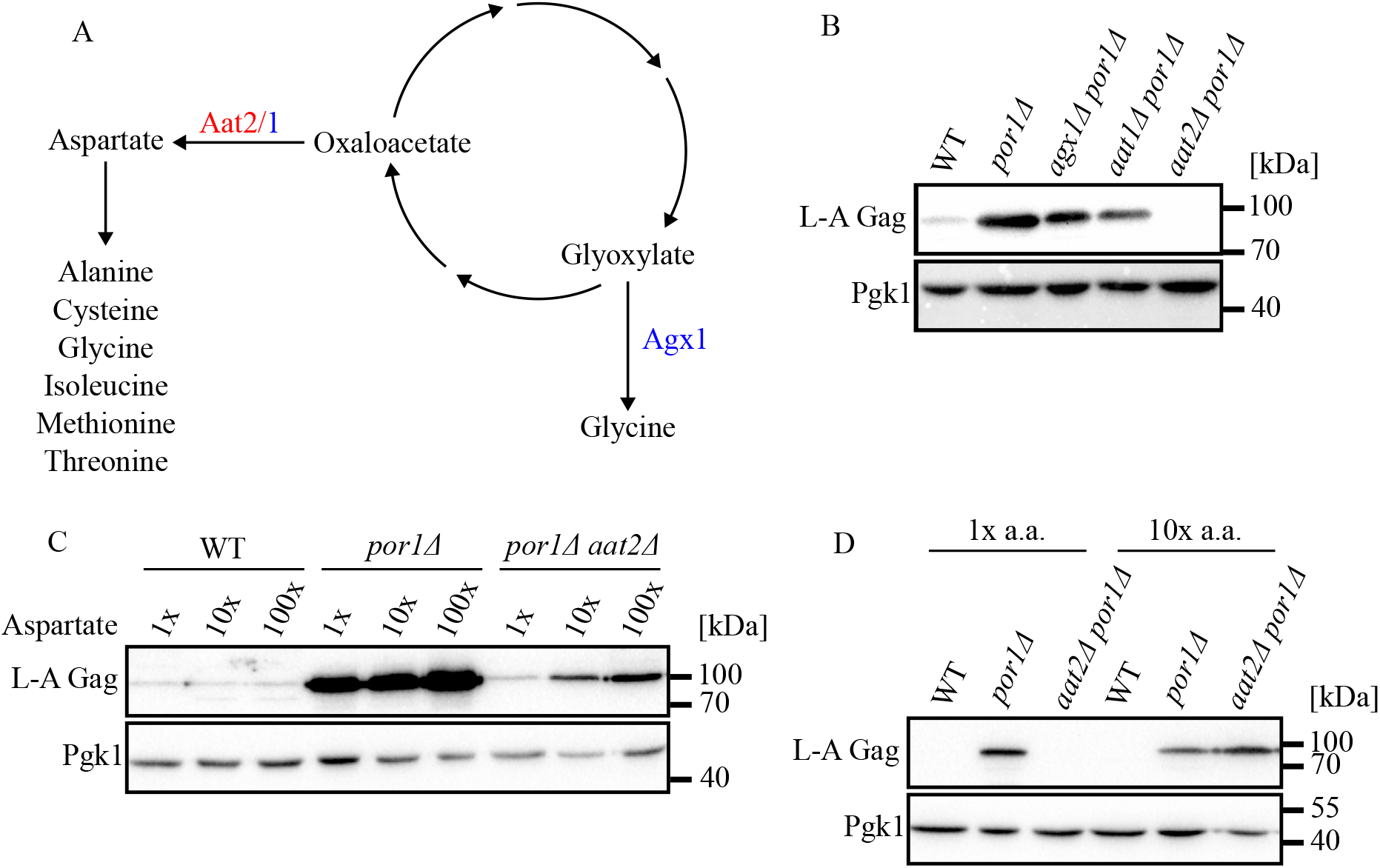
Amino acid synthesis produces the biomolecules necessary for viral replication in a nutrient deplete environment. **(A)** Schematic of amino acids synthesis pathways from the glyoxylate cycle. Glyoxylate is converted into glycine via the alanine:glyoxylate transferase Agx1, and the aspartate aminotransferase Aat1/2 converts oxaloacetate to aspartic acid. Enzymes tested to be required for viral replication are shown in red, while enzymes dispensable for viral replication are shown in blue. **(B-D)** Western blotting of L-A Gag and Pgk1 protein levels in the indicated strains. Samples were collected from 7-day cultures grown in SC media supplemented with glucose. Molecular weight markers are indicated on the right. Additionaly, samples were either supplemented with the indicated amount of **(C)** additional aspartic acid or **(D)** amino acids.

If Aat2 promotes L-A replication in stationary phase by promoting the synthesis of aspartate, then aspartate supplementation in the media might restore high L-A viral load in *por1Δ aat2Δ*. Confirming this prediction, *por1Δ aat2Δ* cultured in increasing amounts of aspartate showed a dose responsive increase in L-A Gag levels (Figure 4C). Since aspartate supplementation alone was insufficient to fully restore Gag levels in *aat2Δ por1Δ*, we tested if supplementing with the full complement of amino acids further enabled L-A replication. Remarkably, we found that *aat2Δ por1Δ* supplemented with 10x amino acids restored Gag accumulation comparable to as in a *por1Δ* single mutant (Figure 4D).

To summarize, here we identify a new yeast antiviral mechanism controlled through Por1/VDAC mediated inhibition of Snf1/AMP Kinase in stationary phase cells. Consistent with many studies showing that Snf1 is activated by glucose starvation, we only observe accumulation of the activated phosphorylated form of Snf1 in post-diauxic phase cells that have consumed all the glucose. Following this transition to glucose starvation, *por1*Δ cells exhibit dramatically increased levels of Snf1-pT210 as well as a 60-fold increase in L-A, suggesting that activated Snf1 promotes L-A replication. Supporting this hypothesis, genetic epistasis experiments show that Snf1 acts with its co-factors and Cat8/Sip4 transcription factor targets to elevate L-A replication in *por1*Δ cells. Through LC-MS/MS, supplementation and genetic experiments, we identify the glyoxylate cycle and amino acid production as the essential outputs of Snf1 signaling that foster L-A replication (Figure 5).

**Figure 5.**
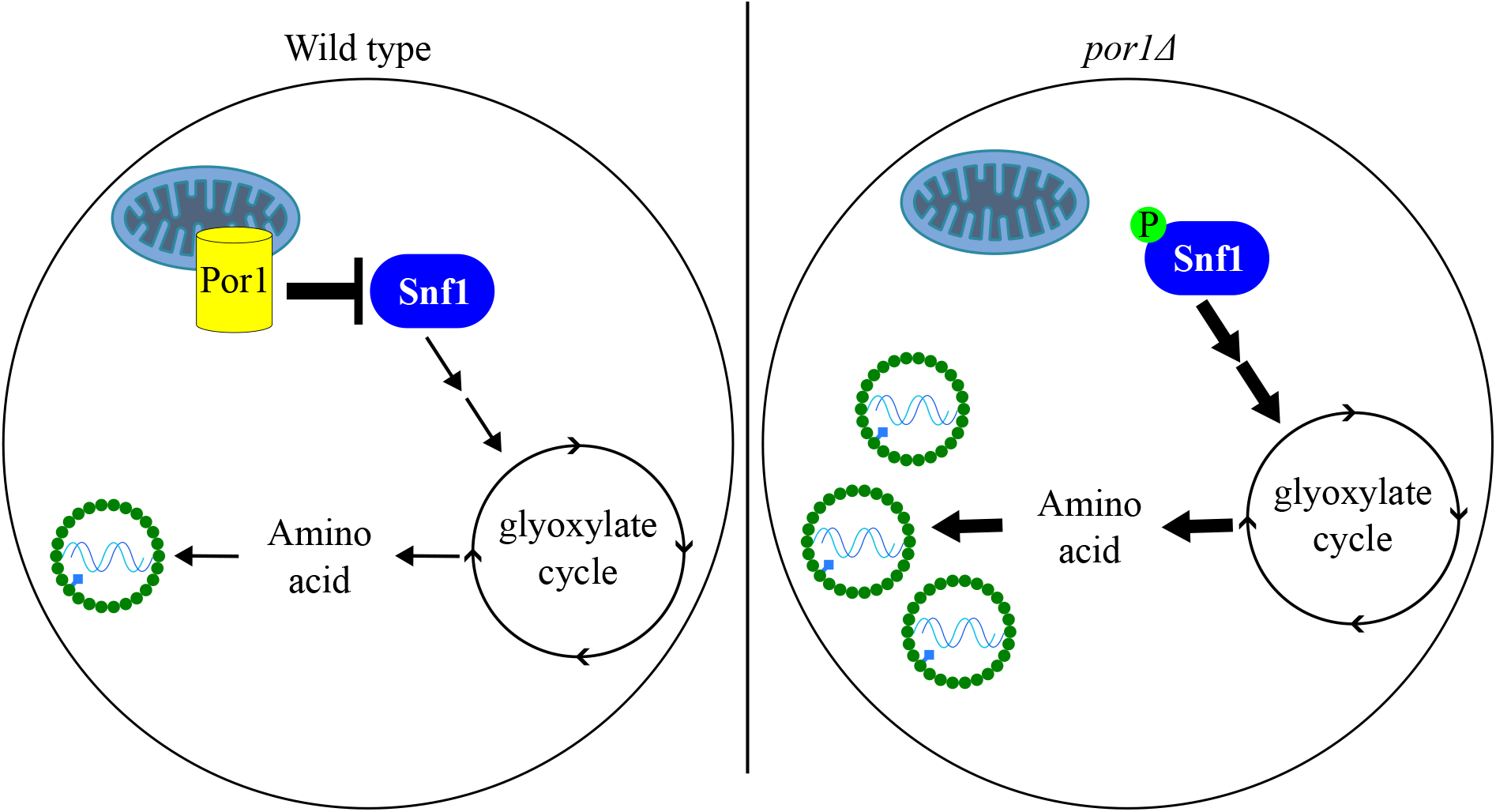
Model of Por1 repression on Snf1 pathway to limit L-A viral replication. In nutritionally starved cells, the AMPK Snf1 is activated to induce the glyoxylate cycle for the synthesis of amino acids. In wild type cells, Por1 negatively represses Snf1 activity to prevent overproduction of amino acid in stationary phase cells. However, when this process is not properly regulated in *por1Δ* strains, the L-A virus hijacks the excess amino acid produced to enhance its replication success.

## Discussion

One of the distinguishing features of mycoviruses is their endemic presence in strains once an infection is established. These infections persist on evolutionary timescales and can adaptively shape fungal physiology raising questions about how these long-term host-virus interactions are mediated in fungi(*2*). Here we identify *POR1*-*SNF1* signaling as a new yeast mechanism that prevents hyper-replication of the L-A totivirus through its control of cellular metabolism. Of note, we only observe *POR1*-*SNF1* signaling in nutrient exhausted stationary phase cells that have exited the cell cycle. In nature, fungal cells likely encounter such conditions frequently, representing a point of vulnerability where mycoviruses could opportunistically replicate in their dormant hosts. Physiological control mechanisms that limit bio-molecules essential for viral replication may thus represent a broadly useful strategy. Indeed, amino acid restriction acts as an antiviral strategy for bacterial phage and human HIV infections(*42, 43*). Given that Snf1 controls allocations towards amino acids after glucose is depleted, any mechanism that can modulate this might provide advantages to the virus during infections(*44*). Despite these connections however, our studies do not necessarily indicate that the sole function of *POR1-SNF1* signaling is antiviral and it is easy to envision other potential roles of this metabolic control system.

A pressing question concerns the molecular mechanisms by which Por1 represses Snf1. Two studies showed Snf1 mitochondrial localization and physical association of Snf1 with Por1 in cells undergoing acute glucose withdrawal(*45, 46*). Although neither study addressed Snf1 activation and signaling such as what we show here, they nevertheless are consistent with a proximal regulatory role of Por1 for Snf1.

Recent findings support a more vivid hypothesis of *POR1-SNF1* signaling and point to Por1’s control of lipid transport/metabolism as the underlying mechanism by which it represses Snf1. Thin layer chromatography and shotgun lipidomics experiments showed that deletion of *POR1* caused myriad defects in bulk phospholipid levels, including reductions in phosphatidylethanolamine (PE) and cardiolipin (CL) along with increased phosphatidic acid (PA) and several others(*23, 25, 47*). Por1 controls these processes in part through its physical interactions with the Ups1/2-Mdm35 intra-mitochondria shuttles that control transport of PA and phosphatidylserine from the outer mitochondrial membrane to the inner membrane(*23*). Notably, these studies investigated dividing cells, and our findings show Por1 function in stationary phase, a context known to involve massive lipid dynamics(*48-50*). A key insight comes from a study of Ups2-Mdm35 in stationary phase cells showing that its mutation caused PE accumulation and was associated with a modest increase in Snf1-pT210 levels, though not to the degree caused by *por1*Δ that we report here(*51*). It was further shown that Snf1 binds to several phospholipids including PA and PE *in vitro* with the implied hypothesis being that Snf1 activity is controlled through sensing of lipids differentially controlled by Ups2-Mdm35(*51*).

All the above-mentioned studies preceded the discovery that Por1 possesses potent lipid scramblase activity in its oligomerized form and how *POR1* influences lipid dynamics in stationary phase remains unknown(*26, 27*). We report here that Por1 levels are strongly increased in stationary phase, which may promote the formation of Por1 oligomers that activate its lipid scramblase activity. An emergent hypothesis is that Por1 induction in stationary phase promotes its oligomerization and scramblase activity, preventing a lipid context that causes Snf1 activation. Pharmacological inhibition of human VDAC has been shown to cause increased Snf1 activity and the human Snf1 homolog AMPK is known to be allosterically activated by long-chain fatty acyl-CoA esters (*52-54*). Evidence from both yeast and human thus suggest an underlying universality of *POR1*-*SNF1* signaling.

## Materials and Methods

### Yeast strains, media, and plasmids

Standard *S. cerevisiae* genetic and strain manipulation techniques were used for strain construction. All strains are derivatives of BY4742 constructed through crossing and dissection. Refer to Table S2 for all yeast strains used. For all experiments, strains were first grown in log phase for at least 10 doublings by serially splitting back of cultures. Time 0 for all stationary phase experiments corresponded to these log phase cells at an OD_600_ of 1. All experiments were carried at 30 °C in synthetic complete media (SC; 0.348% yeast nitrogen base, 1% ammonium sulphate, and 2% glucose) with the appropriate amino acid powder mix (Sunrise Science) unless otherwise specified.

### Measurement of yeast growth and physiological activity

Saturated cultures were diluted to an OD_600_ of 0.1 in 200 µL in a 96-well plate. The plate was sealed a Breathe-Easy membrane (MilliporeSigma) and growth curve data was generated at 30 °C using an S&P growth curve robot (S&P Robotics Inc). Plates were shaken, and the optical density readings were taken every 15 min for 24 h. The data were plotted using R studio ggplot2. Glucose and ethanol concentration in media were measured using assay kits from MyBioSource (MBS8243232 and MBS8309715) according to the manufacturer’s instructions. The luminescent signals were detected using Varioskan LUX Multimode Microplate Reader (ThermoFisher).

### Protein extraction and western blotting

Cells were harvested at indicated time points and permeabilized with 0.1N NaOH at room temperature for 5 min. The cells were then pelleted and resuspended in SDS/PAGE buffer. Cells were disrupted by bead-beating for 3 minutes before heating at 100°C for 10 minutes. The samples were centrifuged to isolate the soluble fraction for western blotting. Protein concentrations were determined with an RC/DC assay (BioRad 5000121). For Snf1-pT210 western blotting, cells were boiled at 100°C for 3 minutes before extraction.

Equal amounts of protein were electrophoresed on 10% SDS-PAGE gels and transferred to polyvinylidene difluoride (PVDF) membranes. Membranes were incubated in primary antibody at 4°C overnight and probed with 1:3,000 horseradish peroxidase (HRP)-conjugated horse anti-mouse (7075; Cell Signaling Technology) or goat anti-rabbit (7074; Cell Signaling Technology) secondary antibody. The proteins were detected with Luminata Forte Western HRP Substrate (EMD Millipore) and imaged with the Bio-Rad ChemiDoc XRS+ system. Images were processed with the Image Lab software package (Bio-Rad). The primary antibodies and their dilutions were 1:1,000 anti-FLAG M2 (F1804; Sigma-Aldrich), 1:1,000 anti-VDAC1/Porin (ab110326; Abcam), 1:5,000 anti-Pgk1 (ab113687; Abcam), 1:2,000 anti–L-A Gag (obtained from Reed Wickner), and 1:1000 anti-Phospho-AMPKα (Thr172) (2535; Cell Signaling Technology).

### RT-qPCR

RNA was prepared from 10 ODs of cells harvested from 7-day cultures and used for RT-qPCR as previously described with some modifications. Briefly, harvested 7-day cell pellets were resuspended in Trizol (15596026; Invitrogen) and subjected to bead-beating (Mini Bead beater, Biospec Products) for 1 minute followed by 2-minute incubations on ice for 8 cycles. Samples were then incubated for 30 minutes at 65°C in acidic phenol (P4682; Sigma-Aldrich), SDS, and buffer AE (10mM Tris-HCl, 0.5mM EDTA pH 9.0) solution. The phase-separated supernatant was washed with chloroform and precipitated overnight. The precipitate was then washed in 70% ethanol and dissolved in water. RNA samples were purified with the RNeasy Mini Kit (74104; Qiagen), and residual DNA was digested with DNase I (79254; Qiagen). 900 nanograms of RNA was reverse transcribed using random nonamers and Maxima H Minus Reverse Transcriptase (EP0753; Thermo Fisher). The cDNA product was isolated by alkaline hydrolysis and treated with RNase A. Subsequently, qPCR was performed on 1/20 dilutions of cDNA product with the SensiFAST SYBR Hi-ROX Kit (BIO-92005; Meridian Bioscience) on the CFX384 platform (BioRad). The data were plotted using R studio ggplot2.

### Metabolite extractions and measurements by LC-MS/MS

Experiment was done as described in (26). Strains were grown overnight in YPAD. Saturated cultures were diluted to 0.02 and 0.05 OD for wild type and *por1Δ*, respectively, and 5 ODs of cells were collected at the 24-hour time point.

## Data availability

Strains are available upon request. The authors affirm that all data necessary for confirming the conclusions of the article are present within the article figures and tables. Supplemental material available at GENETICS online.

## Acknowledgements

We thank Dr. Alex Ensminger for use of the S&P growth curve robot.

---

**Figure S1.**
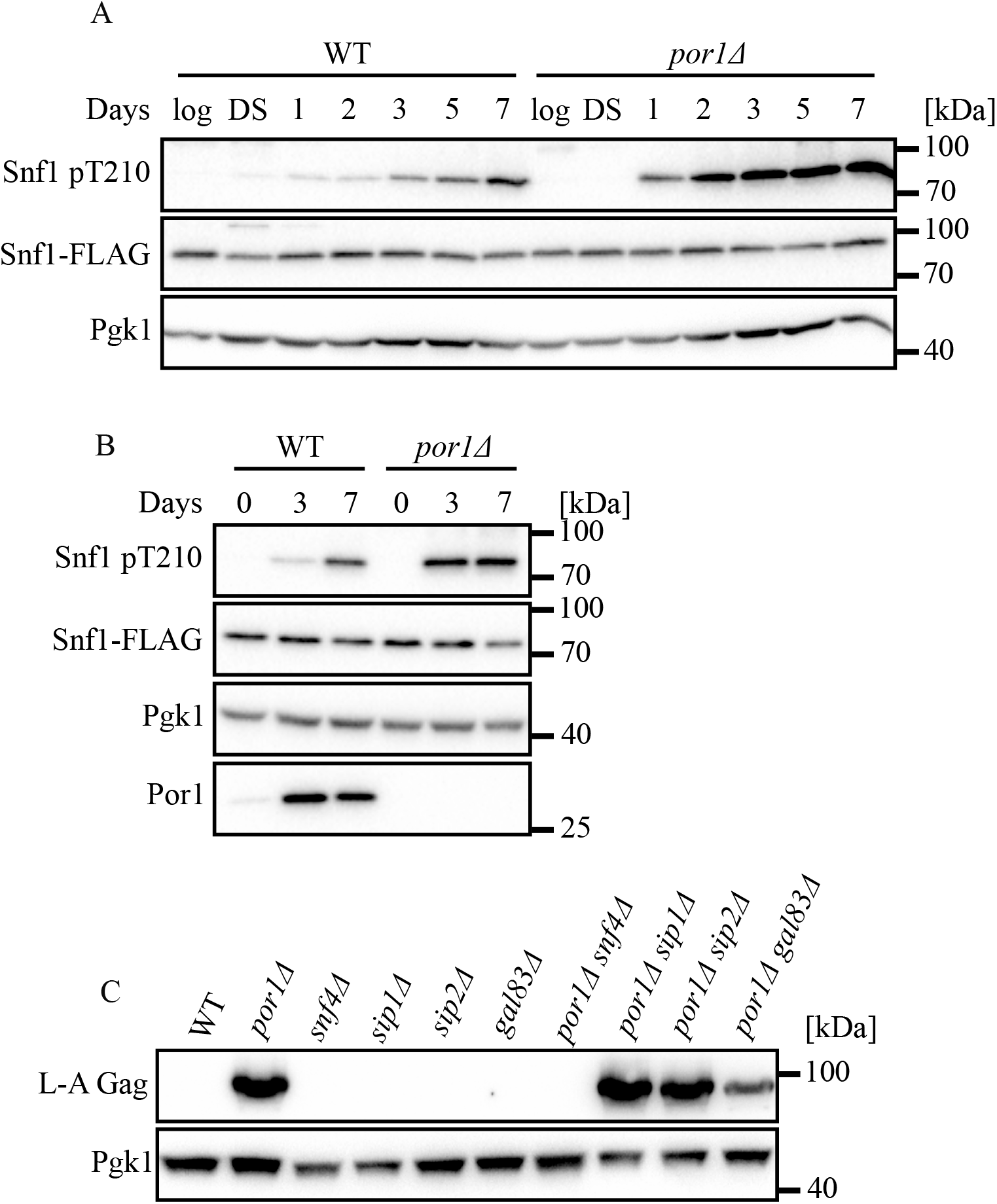
Por1 represses Snf1 activity specifically in stationary phase. **(A, B)** Western blotting of phosphorylated Snf1, Snf1-FLAG, Pgk1, and/or Por1 protein levels in the indicated strains. Samples were collected from cultures grown for the indicated time points in YPAD media. Molecular weight markers are indicated on the right. **(C)** Western blotting of L-A Gag and Pgk1 protein levels in the indicated strains. Samples were collected from 7-day cultures grown in SC media supplemented with glucose. Molecular weight markers are indicated on the right.

**Figure S2.**
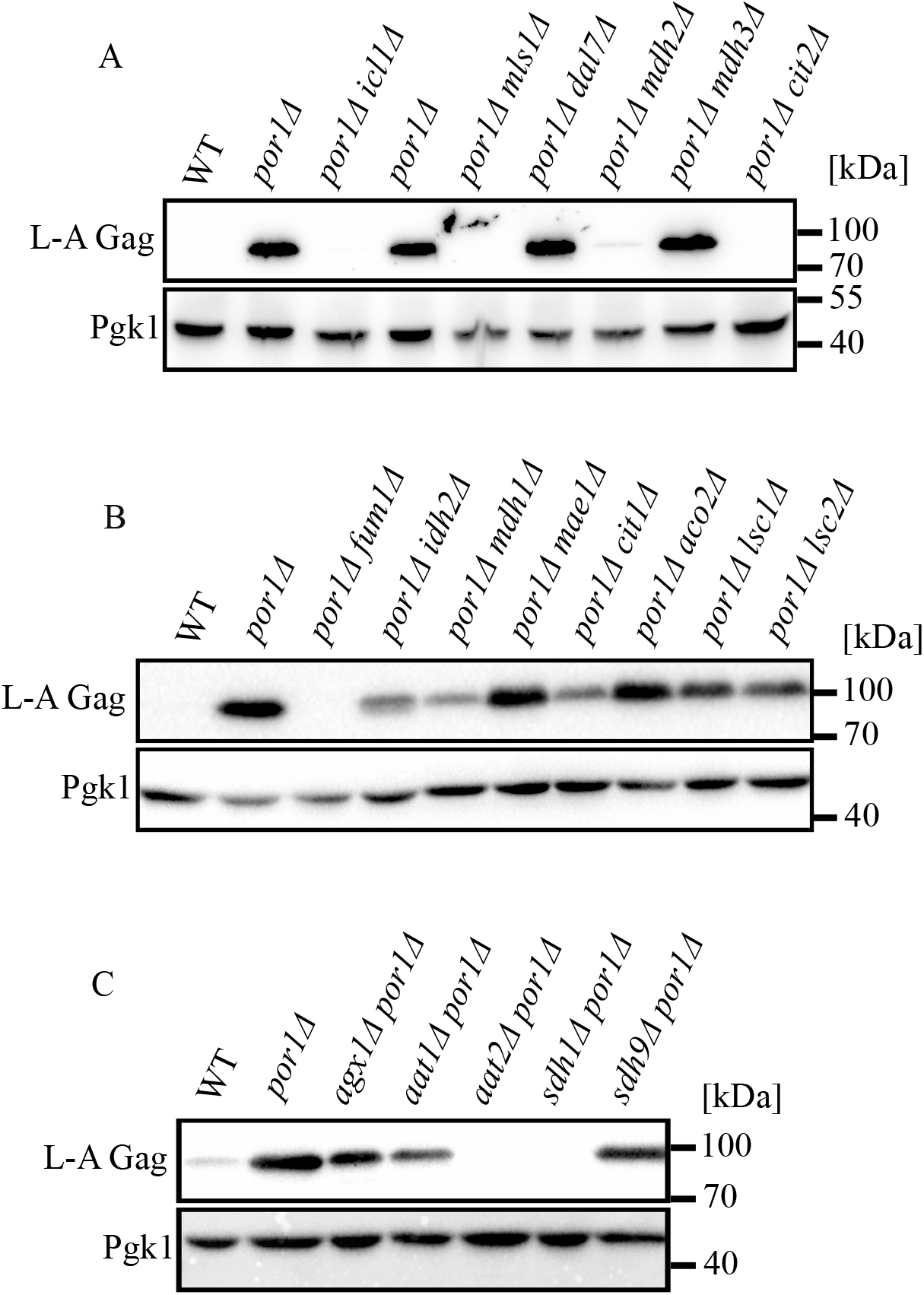
TCA cycle is not required for L-A replication in stationary phase. **(A-C)** Western blotting of L-A Gag and Pgk1 protein levels in the indicated strains. Samples were collected from 7-day cultures grown in SC media supplemented with glucose. Molecular weight markers are indicated on the right.

